# APOBEC3 activity promotes the survival and evolution of drug-tolerant persister cells during acquired resistance to EGFR inhibitors in lung cancer

**DOI:** 10.1101/2023.07.02.547443

**Authors:** Nina Marie G. Garcia, Jessica N. Becerra, Brock J. McKinney, Ashley V. DiMarco, Feinan Wu, Matthew Fitzgibbon, James V. Alvarez

## Abstract

APOBEC mutagenesis is one of the most common endogenous sources of mutations in human cancer and is a major source of genetic intratumor heterogeneity. High levels of APOBEC mutagenesis are associated with poor prognosis and aggressive disease across diverse cancers, but the mechanistic and functional impacts of APOBEC mutagenesis on tumor evolution and therapy resistance remain relatively unexplored. To address this, we investigated the contribution of APOBEC mutagenesis to acquired therapy resistance in a model of EGFR-mutant non-small cell lung cancer. We find that inhibition of EGFR in lung cancer cells leads to a rapid and pronounced induction of APOBEC3 expression and activity. Functionally, APOBEC expression promotes the survival of drug-tolerant persister cells (DTPs) following EGFR inhibition. Constitutive expression of APOBEC3B alters the evolutionary trajectory of acquired resistance to the EGFR inhibitor gefitinib, making it more likely that resistance arises through *de novo* acquisition of the T790M gatekeeper mutation and squamous transdifferentiation during the DTP state. APOBEC3B expression is associated with increased expression of the squamous cell transcription factor ΔNp63 and squamous cell transdifferentiation in gefitinib-resistant cells. Knockout of p63 in gefitinib-resistant cells reduces the expression of the ΔNp63 target genes IL1α/β and sensitizes these cells to the third-generation EGFR inhibitor osimertinib. These results suggest that APOBEC activity promotes acquired resistance by facilitating evolution and transdifferentiation in DTPs, and suggest that approaches to target ΔNp63 in gefitinib-resistant lung cancers may have therapeutic benefit.

## INTRODUCTION

The APOBEC mutational signature, characterized by C-to-T transitions and C-to-G transversions occurring in a TpCpW context, is a prominent mutational process in multiple tumor types, including lung adenocarcinomas^1–5^. APOBEC mutagenesis is due to the activity of the APOBEC3A and APOBEC3B enzymes, which deaminate cytosines on single-stranded DNA^6–9^. APOBEC mutagenesis and expression of *APOBEC3A* and *APOBEC3B* genes are associated with poor prognosis and are increased in metastatic and drug-resistant tumors^10–12^. Therefore elucidating the functional consequences of APOBEC mutagenesis on tumor progression, and the mechanisms underlying these effects, is an important goal.

To address these issues, we examined the role of APOBEC mutagenesis in acquired resistance to EGFR inhibitors in EGFR-mutant lung cancer. First-generation EGFR inhibitors, such as gefitinib and erlotinib, exhibit activity in patients with non-small cell lung cancers (NSCLC) harboring activating mutations in EGFR^13–15^. However, nearly all patients will develop resistance to these targeted therapies, limiting treatment options for these patients^16–18^. Consequently, it is important to identify and characterize mechanisms of acquired drug resistance to EGFR inhibitors. The most common mechanisms of resistance are secondary mutations in the EGFR gene itself, including the T790M mutation, which prevents drug binding and allows persistent EGFR activity even in the presence of drug^19,20^. In this subset of patients, third-generation EGFR inhibitors, such as osimertinib, are often the next treatment option^21^. A number of other genetic resistance mechanisms have been discovered, including amplification or mutation of other receptor tyrosine kinases, including Met, FGFR , HER2, and ALK^22–25^. Several non-genetic resistance mechanisms have also been described, including the epithelial-to-mesenchymal transition (EMT)^26–28^ and histological transformation to other lung cancer cell types, such as small-cell lung cancer (SCLC) and squamous cell carcinoma (SCC)^29–31^. The cellular and molecular pathways underlying histological transformation remain unknown, and importantly there are currently no therapies available for patients who develop resistance to EGFR inhibitors via transdifferentiation to a small-cell or squamous cell phenotype.

Interestingly, the APOBEC mutational signature is elevated in tumors from patients who have undergone treatment with EGFR inhibitors and developed acquired resistance to these inhibitors^32^. Among these patients, APOBEC signatures are especially high in tumors that have acquired resistance through histological transformation^33,34^. Recent work has found that APOBEC3A and APOBEC3B genes are induced by anti-EGFR targeted therapy and contribute to the acquisition of drug resistance^35,36^. However, the precise mechanism by which APOBEC3 enzymes promote resistance to EGFR inhibitors remains unclear. In addition, the relationship between APOBEC mutagenesis and transdifferentiation remains unexplored. In the current study, we explore the regulation and function of APOBEC mutagenesis during acquired resistance to EGFR inhibitors. We find that EGFR inhibition induces APOBEC3 expression and activity, which in turn promotes the survival of DTPs. Sustained APOBEC3B expression promotes the evolution of drug resistance in DTPs and is associated with squamous cell transdifferentiation in EGFR inhibitor-resistant cells. These findings reveal an important role for APOBEC activity in the evolution of drug resistance.

## MATERIALS AND METHODS

### Tissue culture and reagents

#### Cell lines

PC9, HCC827, and BT474 cells were grown in RPMI-1640 media containing 10% fetal bovine serum, L-glutamine, and penicillin/streptomycin. SKBR3 cells were grown in DMEM media containing 10% fetal bovine serum, L-glutamine, and penicillin/streptomycin. Lentivirus from HEK293T cells was produced as previously described^37^. Cells were selected for using puromycin or blasticidin, as indicated.

#### Generation of Cre-inducible A3B expression in PC9 cells

To create a Cre-inducible A3B construct, we used an approach based upon the XTR system^38^. Briefly, the human A3B coding sequence (NM_004900.4) with a C-terminal HA tag and an SV40 intron was cloned downstream of the human PGK promoter in the reverse orientation. This reverse A3B-HA cDNA was flanked by two pairs of incompatible loxP sites (Lox5171 and Lox2722), such that expression of Cre recombinase leads to inversion of the A3B cDNA and expression of the full-length A3B gene. This construct, pLenti-APOBEC3B-Intron-HA-Cre Flp, was generated by VectorBuilder.

#### CRISPR/Cas9-mediated knockout

lentiCas9-Blast was a gift from Feng Zhang (Addgene plasmid # 52962). For A3A and A3B knockout, PC9 Cas9 cells were transduced with lentiviral constructs expressing dual sgRNAs, either non-targeting (NT1/NT2), A3A/NT1, A3B/NT1, or A3A/A3B. NT1 gRNA: GGCAGTCGTTCGGTTGATAT; NT2 gRNA: GCTTGAGCACATACGCGAAT; A3A gRNA: GTGCTGGTCCATCTTGACCG; A3B gRNA:

ATGACCCTTTGGTCCTTCGA. Dual-sgRNA vectors were generated by Cellecta. Knockout cells were selected using puromycin and then transduced with H2B-GFP lentivirus. GFP-positive knockout cells were sorted for using flow cytometry and single clones were generated.

For TP63 knockout, gRNAs targeting TP63 were cloned into lentiCRISPR v2 (a gift from Feng Zhang, Addgene plasmid # 52961). An empty LCV2-Cas9 construct was used as a control. TP63 sgRNA 1: CAATGATTAAAATTGGACGG. TP63 sgRNA 3: GCTGAGCCGTGAATTCAACG.

#### A3B expression and drug treatments

For expression of A3B, 2×10^5^ PC9 Cre/Flp A3B cells were seeded onto a 6-well plate. 24 hr later, adenovirus expressing Cre recombinase (Vector Biolabs) was added to the media at an MOI of 1000. For short-term treatments, gefitinib (SelleckChem) was used at a concentration of 500 nM and osimertinib (SelleckChem) was used at a concentration of 1 µM. For long-term cell viability assays, PC9 cells were treated with 100 nM osimertinib.

#### Knockdown experiments

SMARTpool ON-TARGETplus siRNAs for RELA, RELB, c-REL, IRF3, and STAT1 were purchased from Horizon. The DharmaFECT transfection protocol was followed as described. Briefly, PC9 cells were plated with antibiotic-free complete media and incubated overnight. After 24 hr, siRNA SMARTpools were resuspended and diluted to the desired concentration. They were then mixed with the transfection reagent (DharmaFECT) in serum-free media and brought up to volume with complete media. The transfection medium was added to the cells. The cells were then incubated with the transfection media for 24 hr. PC9 cells were treated with gefitinib or osimertinib 48 hr after transfection. Cells were harvested for subsequent experiments 24 hr after drug treatment.

#### Resistance assays

2×10^5^ PC9 Cre/Flp A3B cells were seeded onto 6-well plates. After 24 hr, AdenoCre recombinase (Vector Biolabs) was added to the media at an MOI of 1000. Cells were cultured for 3 weeks prior to treatment with gefitinib. After 3 weeks, 1×10^6^ PC9 cells, with or without A3B expression, were seeded onto 10 cm plates. 24 hr later, media was changed to media containing 100 nM gefitinib. Media was changed every 3 days post-treatment and cells were counted and passaged every 7 days post-treatment until cells acquired resistance. 1×10^6^ cells (or the entire culture if less than 1×10^6^ cells) were replated at each passage. Gefitinib-resistant cells were cultured continuously in gefitinib-containing media. Calculated cell number was determined using the following formula:

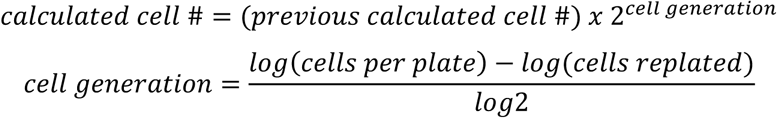

### qRT-PCR

RNA was isolated from cells using RNeasy columns (Qiagen). 1 μg of RNA was reversed transcribed using cDNA synthesis reagents (Promega). qPCR was performed using the following 6-carboxyfluorescein-labeled TaqMan probes (Thermo): APOBEC3A (Hs00377444_m1 ), APOBEC3B (Hs00358981_m1), ACTB (Hs01060665_g1), 18S (Hs03003631_g1) IL1A (Hs00174092_m1), IL1B (Hs01555410_m1), dN-TP63 (Hs00978339_m1), total TP63 (Hs00978340_m1), RELA (Hs01042014_m1), RELB (Hs00232399), C-REL (Hs00968440_m1), IRF3 (Hs01547283_m1), STAT1 (Hs01013996_m1). qPCR was performed using an AB Applied Biosystems ViiA7 qPCR machine.

### Western Blotting

Western blotting was performed as described^37^ using the following antibodies: APOBEC3A/B/G^9^ (gift from Maciejowski Lab), actin (Cell Signaling Technology), HA-tag (Cell Signaling Technology), and α-tubulin (Cell Signaling Technology), all at a 1:1000 dilution. Secondary antibodies conjugated to Alexa Flour 680 (Life Technologies, Carlsbad, CA) or IR-800 (LI-COR Biosciences, Lincoln, NE) were detected with the Odyssey detection system (LI-COR Biosciences). For endogenous A3B detection, secondary antibodies conjugated to HRP were used and blots were developed using Forte or Crescendo reagent (Millipore, Burlington, MA) and exposed to film (VWR, Radnor, PA). Secondary antibodies were used at a 1:5000 dilution.

### Deaminase assay

Deaminase assays were performed as previously described^37^. % deamination was calculated using the following formula:

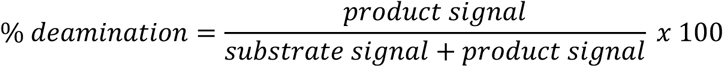

### ddPCR

Genomic DNA was isolated using the DNeasy Blood and Tissue Kit (Qiagen). Each reaction contained the following: 10ng gDNA, EGFR T790M probe (Assay ID: dHsaCP2000019, BioRad), wild-type EGFR probe (Assay ID: dHsaCP2000020, BioRad), HaeIII restriction enzyme (New England Biolabs), and ddPCR Supermix no dUTPs (BioRad). 20 µl of each reaction was loaded onto a DG8 cartridge (BioRad), along with 70 µl of droplet generation oil (BioRad). Droplets were generated using the BioRad QX200 Droplet Digital PCR System and then transferred to a 96-well plate for PCR. PCR conditions were as follows: 95°C for 10 min, 40 cycles of 94°C for 30 sec followed by 55°C for 1 min, and 98°C for 1 min. Droplets were analyzed on the BioRad QX200 Droplet Reader for FAM (T790M) and HEX (WT) probes. Data was analyzed with the QuantaSoft software (BioRad) to obtain fractional abundance of mutated gDNA. % EGFR T790M was calculated using the following formula:

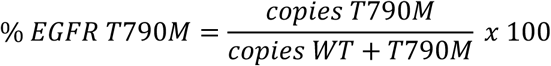

### Competition assay and flow cytometry

GFP-labeled A3A/A3B double knockout clones were mixed with NT1/NT2 control cells in a 1:1 ratio and then seeded onto 6-well plates. After a 24 hr, cells were treated with either 100 nM gefitinitib or 500 nM osimertinib. Media containing drug or vehicle was changed every 3-4 days. On days 7 and 14 after drug treatment, the ratio of GFP-positive to GFP-negative cells was measured using flow cytometry on a BD FACSCanto II flow cytometer.

### RNA sequencing

RNA was isolated from cells using RNeasy columns (Qiagen). RNA-sequencing was performed by Novogene. Briefly, mRNA was purified from total RNA using poly-T oligo-attached magnetic beads. After fragmentation, the first strand cDNA was synthesized using random hexamer primers, followed by the second strand cDNA synthesis using dTTP for non-directional library. Libraries were subjected to end-repair, A-tailing, adapter ligation, size selection, amplification, and purification. Quantified libraries were pooled and sequenced on NovaSeq PE150.

STAR v2.7.9a with 2-pass mapping was used to align paired-end reads to human genome assembly hg38 and GENCODE gene annotation v38 along with gene-level read quantification. Bioconductor package edgeR 3.36.0 was used to detect differential gene expression between sample groups. Protein-coding and lncRNA genes were included in the analysis. Genes with low expression were excluded using edgeR function filterByExpr with min.count = 10 and min.total.count = 15. The filtered expression matrix was normalized by TMM method and subject to significance testing using quasi-likelihood pipeline implemented in edgeR. For group comparisons, biological replicates are used as blocking factor (i.e. unwanted covariate) in the statistical model when applicable. A gene was deemed differentially expressed if absolute log2 fold change was above 1 (i.e. fold change > 2 in either direction) and Benjamini-Hochberg adjusted p-values was less than 0.01.

### Whole exome sequencing

Genomic DNA was isolated using the DNeasy Blood and Tissue Kit (Qiagen). Twist Biosciences WES Kit was used for capture and library prep and paired-end Illumina next-generation sequencing was performed using NextSeq2 according to manufacturer’s recommendations. Read processing and germline variant calling followed GATK best practice workflow (https://gatk.broadinstitute.org/hc/en-us/articles/360035535932-Germline-short-variant-discovery-SNPs-Indels-). Known SNVs and INDELs were retrieved from GATK resource bundle (https://console.cloud.google.com/storage/browser/genomics-public-data/resources/broad/hg38/v0;tab=objects?prefix=&forceOnObjectsSortingFiltering=false). Briefly, Illumina adapters were first trimmed from paired-end reads using cutadapt (http://dx.doi.org/10.14806/ej.17.1.200) and trimmed reads were mapped to reference genome hg38 using BWA 0.7.17 (https://doi.org/10.1093/bioinformatics/btp324). Bam files were further processed using GATK 4.2.5 to generate analysis-ready reads that have proper read group information, duplicated reads marked (MarkDuplicates), and recalibrated base quality score (BaseRcalibrator and ApplyBQSR). GATK HaplotypeCaller was run in GVCF mode for each sample followed by joint calling of all samples using GenomicsDBImport and GenotypeGVCFs. Twist panel targeted regions were padded with 50bp on both sides for the above processing and variant calling. Due to less than 30 samples in the cohort, Variants were subject to hard filtering by following the GATK article (https://gatk.broadinstitute.org/hc/en-us/articles/360035531112--How-to-Filter-variants-either-with-VQSR-or-by-hard-filtering#1).

### Xenograft studies

1×10^5^ cells diluted in PBS:Matrigel (1:1) were injected subcutaneously into the flanks of nude mice (6-8 weeks old). Tumors developed in 6-8 weeks and tumor volume was measured once a week using calipers. Established tumors were harvested and fixed in 10% neutral buffered formalin for 24 hr, washed in PBS, and then stored in 70% ethanol.

### Immunohistochemistry

Staining was performed by the Experimental Histopathology core resource at Fred Hutchinson Cancer Center. For cell lines, 5 × 10^6^ cells were harvested, washed in PBS, and fixed in 200 µl 10% neutral buffered formalin for 24 hr. Tumors were harvested and fixed in 10% neutral buffered formalin for 24 hr, washed with PBS, and then stored in 70% ethanol until staining. Tumor sections and cells were stained with p40 antibody (Abcam) to detect ΔNp63.

### Statistics

Statistical analysis was performed using GraphPad Prism. Statistical tests used and p-values are reported with each figure.

## RESULTS

### APOBEC3 expression and activity are induced following EGFR inhibition

To explore the functional consequences of APOBEC mutagenesis on acquired resistance to EGFR inhibitors we used PC9 cells, a NSCLC cell line that has an activating mutation in EGFR (exon 19 deletion). We first assessed how the expression and activity of APOBEC3 genes changes in response to EGFR inhibition. Treatment of PC9 cells with the tyrosine kinase inhibitors (TKIs) gefitinib or osimertinib led to induction of both the *APOBEC3A* (A3A) and *APOBEC3B* (A3B) genes (Fig. 1A). In a second EGFR-mutant NSCLC cell line, HCC827, A3A and A3B genes were also induced following treatment with gefitinib and osimertinib (Fig. S1A). A3B protein levels also increased following treatment with these inhibitors (Fig. 1B), though we were unable to detect A3A protein (data not shown). To assess how EGFR inhibition affects the enzymatic activity of APOBEC3 proteins, we used a deaminase activity assay that measures the ability of a lysate to deaminate a cytosine within an APOBEC consensus sequence in a single-stranded DNA oligonucleotide ^39^. Gefitinib or osimertinib treatment led to a 2- to 3-fold increase in deaminase activity (Fig. 1C – D). Together these results suggest that EGFR inhibition in NSCLC cells leads to the induction of APOBEC expression and activity, consistent with previous findings^35,36^.

**Figure 1.**
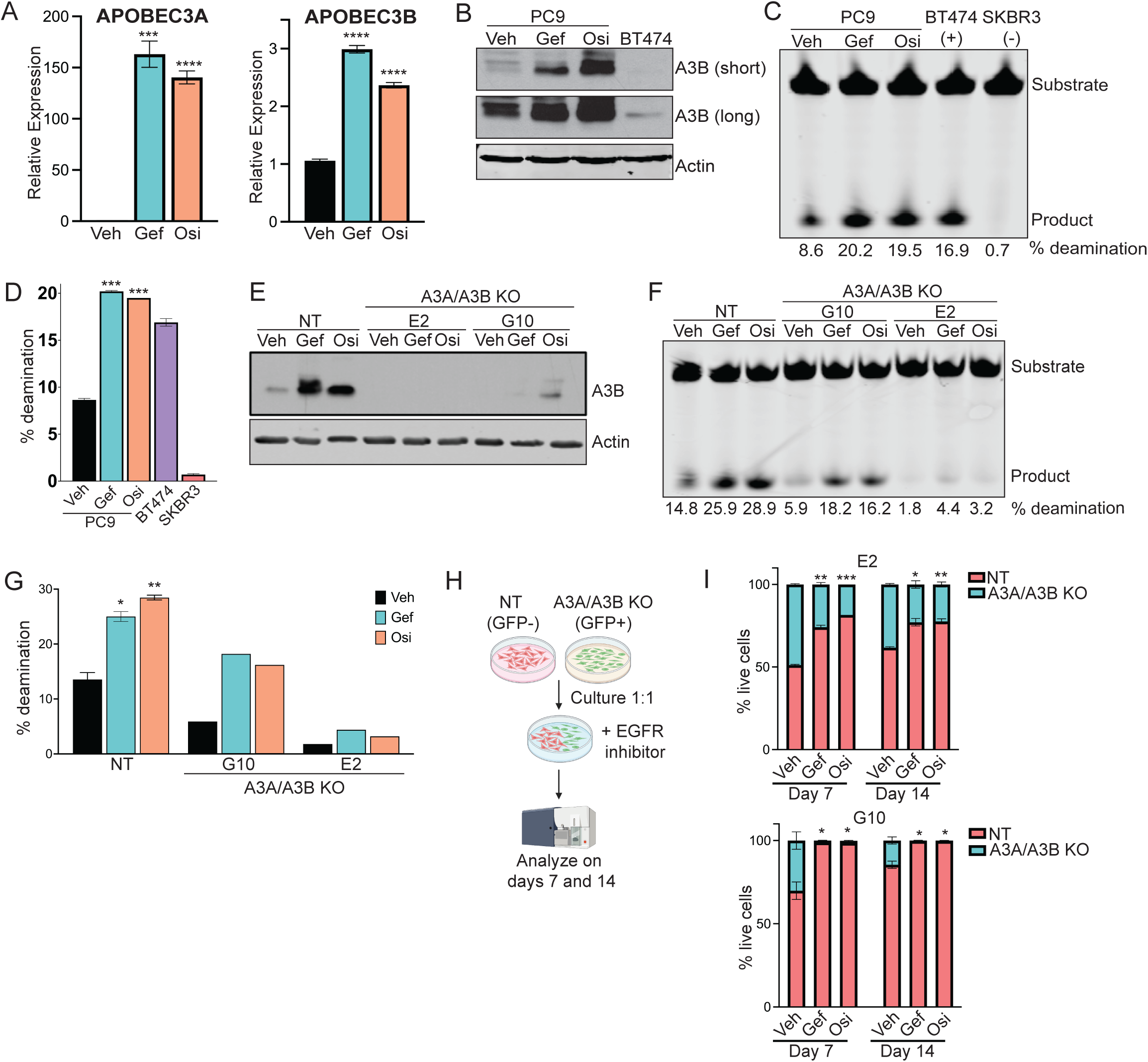
APOBEC3 activity is induced following EGFR inhibition. A. qRT-PCR analysis of a representative experiment showing A3A and A3B expression in PC9 cells following treatment with gefitinib or osimertinib for 24 hours. Error bars represent SEM of two biological replicates. Significance relative to the vehicle was determined using unpaired t-test. *** indicates p<0.0005 and **** indicates p<0.0001. B. Western blot for A3B protein in PC9 cells following treatment with gefitinib or osimertinib for 24 hours. BT474 cells are shown as a positive control. Short and long exposures for A3B are shown. C. In vitro deaminase activity of PC9 cells treated with gefitinib or osimertinib for 24 hours. BT474 and SKBR3 cells are shown as positive and negative controls, respectively. % deamination was calculated as described in Methods. D. Quantification of % deamination shown in C. Error bars represent SEM of two technical replicates. Error bars represent SEM of two technical replicates. Significance relative to the vehicle was determined using unpaired t-test. *** indicates p<0.0005. E. Western blot for A3B protein in PC9 cells expressing non-targeting gRNAs (NT) or gRNAs targeting both A3A and A3B (A3A/A3B KO). A pooled population is shown for NT cells, and two clones are shown for A3A/A3B KO cells. Cells were treated with gefitinib or osimertinib for 24 hours. F. In vitro deaminase activity of PC9 cells expressing non-targeting gRNAs (NT) or gRNAs targeting both A3A and A3B (A3A/A3B KO). A pooled population is shown for NT cells, and two clones are shown A3A/A3B KO cells. Cells were treated with gefitinib or osimertinib for 24 hours and % deamination was calculated as described in Methods. G. Quantification of % deamination shown in F. Error bars represent SEM of two technical replicates. Significance relative to the NT vehicle was determined using unpaired t-test. * indicates p<0.05 and ** indicates p<0.005. H. Schematic of competition assay via flow cytometry. PC9 cells expressing non-targeting gRNAs (NT) or gRNAs targeting both A3A and A3B (A3A/A3B KO) were used for the experiment. I. Quantification of % live cells, either GFP-negative or -positive, after 7 or 14 days of EGFR inhibition. A pooled population is shown for NT cells and two clones are shown for A3A/A3B KO cells. Error bars represent SEM of two biological replicates. Unpaired t-test was used to determine statistical significance o2f 5GFP-positive cells in treated condition relative to vehicle. * indicates p<0.05, ** indicates p<0.005 and *** indicates p<0.0005.

To investigate the pathways responsible for APOBEC3 induction, we used siRNA to knock down candidate transcription factors known to play a role in APOBEC3 expression. We first focused on the NFκB pathway, since these proteins have been shown to regulate APOBEC3 expression in diverse contexts^40–42^. We found that knockdown of RelA did not affect the induction of A3A or A3B (Fig. S1B), while knockdown of RelB or c-Rel partially blunted the upregulation of A3A and A3B following gefitinib or osimertinib treatment (Fig. S1C – D). However, knockdown of these proteins only partially blocked A3A and A3B induction, suggesting that other pathways may contribute to the transcriptional upregulation of APOBEC3 in response to EGFR inhibition.

Other pathways that can activate A3A and A3B include the type-I interferon (IFN) pathway, which signals through STAT transcription factors, and IRF3, which is activated as part of the innate immune response to viral infection^43–45^. We therefore tested whether knockdown of STAT1 or IRF3 prevented A3A and A3B induction following EGFR inhibition. We found that STAT1 knockdown almost completely blocked the upregulation of A3A and A3B following EGFR inhibition (Fig. S1E). In contrast, IRF3 blocked the upregulation of A3B, but not A3A (Fig. S1F). Taken together, these data identify several pro-inflammatory transcription factors, notably STAT1, that are required for the transcriptional upregulation of A3A and A3B following EGFR inhibition.

### APOBEC3 expression promotes the survival of drug-tolerant persister cells following EGFR inhibition

To assess how A3A and A3B upregulation affects the response of lung cancer cells to EGFR inhibition, we used CRISPR-Cas9 to knock out each gene in PC9 cells. Cells were infected with lentivirus expressing Cas9 along with sgRNAs targeting A3A, A3B, or both, and following selection individual clones were expanded and screened. Western blot analysis confirmed that two A3B-knockout clones (B6 and F11) had reduced levels of A3B expression, both at baseline and following EGFR inhibition (Fig. S2A). Because we could not detect A3A protein, we assessed A3A knockout by sequencing genomic DNA surrounding the sgRNA recognition site. Two A3A-knockout clones (B10 and D3) had mutations predicted to disrupt A3A expression (Supplemental Table 1). Using similar approaches, we identified two A3A/A3B-double knockout clones (G10 and E2) with knockout of both A3A and A3B (Fig. 1E and Supplemental Table 1).

We first examined the induction of cytosine deaminase activity in these clones following EGFR inhibition. Knockout of A3B alone, as well as knockout of A3A and A3B together, led to a reduction in basal deaminase activity and blunted the increase in deaminase activity resulting from EGFR inhibition (Fig. 1F – G, Fig. S2B – C). In contrast, knockout of A3A alone had no effect on basal or induced deaminase activity. This suggests that A3B is largely responsible for the increased deaminase activity in PC9 cells following EGFR inhibition.

We next used a cellular competition assay to assess the functional effects of A3A/A3B knockout on the response of PC9 cells to EGFR inhibition. While the majority of PC9 cells die in response to EGFR inhibition, a small population of cells survives treatment and persists as quiescent or slow-growing drug-tolerant persister cells (DTPs)^46–48^. Cells in the DTP state can undergo continued evolution to become fully drug resistant^47^, and pathways that regulate DTP survival can promote or forestall therapy resistance^46^. We therefore tested whether the induction of A3A and A3B following EGFR inhibition regulates the survival of DTPs. Control PC9 cells expressing a non-targeting sgRNA were mixed in a 1:1 ratio with GFP-labeled, A3A/A3B double-knockout clones. Cells were then treated with vehicle, gefitinib, or osimertinib, and the proportion of GFP-positive cells was measured by flow cytometry after 7 and 14 days (Fig. 1H). In the absence of EGFR inhibitors, control and A3A/A3B knockout cells were present at approximately equal proportions, though clone G10 had a slight competitive disadvantage (Fig. 1I). In contrast, A3A/A3B double-knockout cells were markedly depleted starting just 7 days following EGFR inhibition and continuing through day 14 (Figure 1I). These results suggest that the induction of A3A and A3B expression promotes the survival of DTPs following EGFR inhibition, corroborating previous findings^35,36^.

### Engineering an inducible system for APOBEC3B expression in PC9 cells

APOBEC mutagenesis is thought to occur in episodic bursts^49^, and A3A and A3B expression in response to inflammatory stimuli is likewise transient^43^. We therefore evaluated how A3A and A3B expression changed during prolonged EGFR inhibition. A3A expression oscillated over 14 days (Fig. 2A). In contrast, A3B expression returned to baseline levels as early as two days following treatment with EGFR TKIs (Fig. 2A). Therefore, we next wished to investigate the effects of sustained APOBEC activity on acquired resistance to EGFR inhibitors. To achieve this, we generated and characterized PC9 cells with a Cre recombinase-inducible A3B construct (Fig. 2B). Infection with adenovirus expressing Cre led to a ∼3-fold increase in A3B mRNA and protein expression (Fig. 2C – D) and a 2-fold increase in deaminase activity (Fig. 2E – 2F). Importantly, this magnitude of increase in A3B expression and deaminase activity was similar to that observed following EGFR inhibition (cf. Figure 1). Overexpression of A3B did not affect the growth or viability of PC9 cells in vitro (Fig. 2G). This system thus offers an opportunity to address the effects of sustained APOBEC3 activity on acquired therapy resistance.

**Figure 2.**
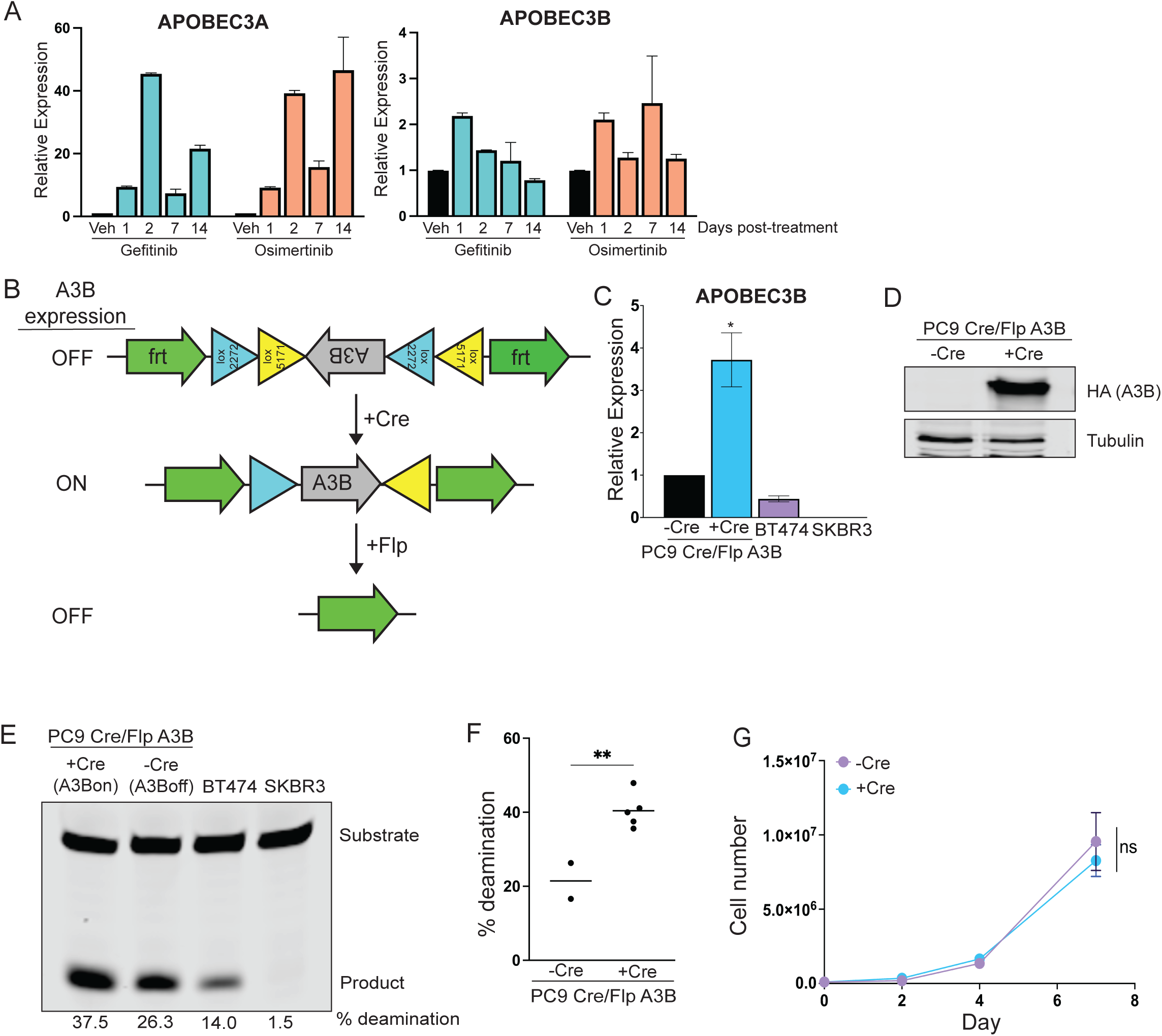
Engineering an inducible system for APOBEC3B expression in PC9 cells. A. qRT-PCR analysis showing A3A and A3B expression in PC9 cells following treatment with gefitinib or osimertinib over the course of 14 days. Error bars represent SEM of three technical replicates. B. Schematic of Cre-inducible APOBEC3B expression. C. qRT-PCR analysis showing A3B expression in PC9 cells following infection with Cre recombinase. Error bars represent SEM of three technical replicates. * indicates p<0.05. BT474 and SKBR3 cells are shown as controls. D. Western blot showing protein expression of HA-tagged A3B in PC9 cells following infection with Cre recombinase. E. In vitro deaminase activity assay in PC9 cells following infection with Cre recombinase. BT474 and SKBR3 cells are shown as controls. % deamination was calculated as described in Methods. F. Quantification of % deamination in two replicates of control PC9 cells (-Cre) and five replicates of A3B-expressing PC9 (+Cre). % deamination was calculated as described in Methods. Unpaired t-test was performed to determine statistical significance. ** indicates p<0.005. G. Growth curves for PC9 cells expressing A3B (+Cre) or control cells (-Cre). Error bars represent SEM of two biological replicates. Two-way ANOVA was performed to determine statistical significance. ns = not significant

### APOBEC3B expression alters the evolutionary trajectory of acquired resistance to EGFR inhibitors

Using PC9 cells with conditional A3B expression, we examined the consequences of sustained A3B expression on acquired resistance to EGFR inhibitors. Five independent replicates of control cells (infected with GFP-expressing adenovirus; denoted as A3B-off A-E) and six independent replicates of A3B-expressing cells (Cre-infected; denoted as A3B-on A-F) were treated with 100 nM gefitinib, and cells were passaged weekly until cells grew consistently in the presence of drug. Aliquots of each replicate cell population not treated with gefitinib, denoted as gefitinib-sensitive, were saved for subsequent comparisons. Consistent with previous studies^47,50^, control PC9 cells evolved resistance to gefitinib over 1-4 months, with one subset of cells developing resistance at around 30–40 days and the other subset developing resistance at later timepoints ranging from 60–100 days (Fig. 3A). A3B-expressing cells developed resistance with the same kinetics as control cells, suggesting that sustained A3B expression does not alter the kinetics with which PC9 cells evolve resistance to gefitinib.

**Figure 3.**
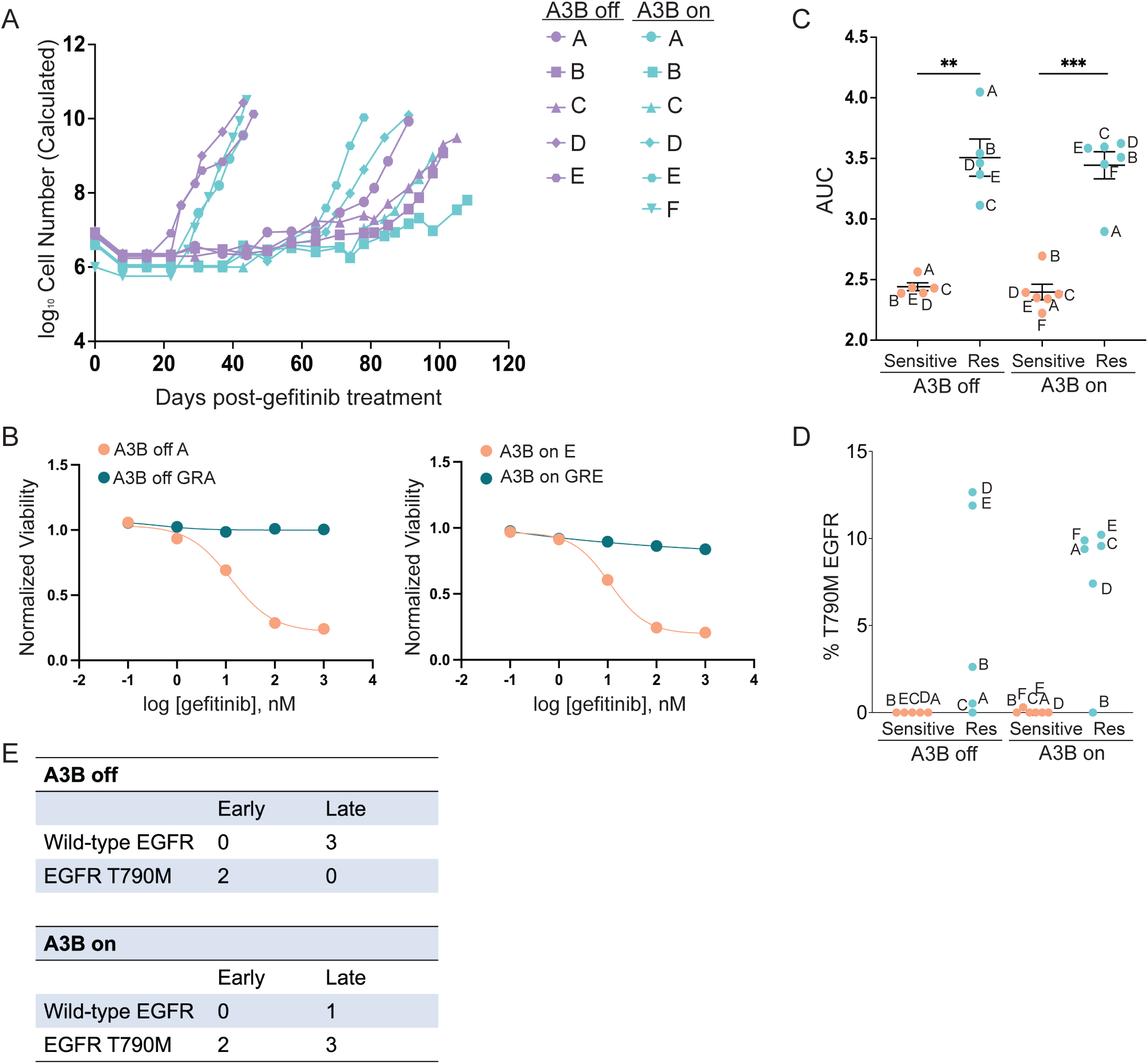
APOBEC3B expression alters the evolutionary trajectory of acquired resistance to EGFR inhibitors. A. Kinetics of evolution of gefitinib resistance in PC9 cells with or without A3B expression. B. Representative dose-response curves to gefitinib for sensitive and resistant A3B-off and A3B-on PC9 cells. C. Quantification of area under the curve (AUC) of dose-response curves to gefitinib for sensitive and resistant A3B-off and A3B-on PC9 cells. Unpaired t-test was performed to determine statistical significance. ** indicates p<0.05. *** indicates p<0.0005. D. Frequency of EGFR T790M mutation as determined by ddPCR in sensitive and gefitinib resistant A3B-off and A3B-on PC9 cells. % EGFR T790M was calculated as described in Methods. E. Contingency table summarizing the relationship between resistance kinetics and E_2_G_7_ FR mutation status in A3B-off and A3B-on PC9 cells.

To further characterize each gefitinib-resistant (GR) control and A3B-expressing population, we measured the concentration of gefitinib required to inhibit growth by 50% (GI_50_) in both sensitive and gefitinib-resistant cells (Fig. 3B, Fig. S3A). The GI_50_ for gefitinib in sensitive cells ranged from 10-20 nM, as expected^47^. In contrast, concentrations of gefitinib up to 1 μM did not affect cell growth in GR cells. We calculated the area under the curve (AUC) for the dose-response curves in sensitive and GR cells. As expected, in both A3B-off and A3B-on contexts resistant cells had greater AUC values, confirming that GR cells are markedly resistant to gefitinib (Fig. 3C). By RNA sequencing analysis, we confirmed that A3B expression remains high in the GR cells (Fig. S5A).

We repeated this resistance assay with an additional set of four control (A3B-off G, I-K) and five A3B-expressing cells (A3B-on G-K) (Fig. S4A). Similar to the previous experiment, we observed that PC9 cells, regardless of A3B expression, acquired resistance to a low dose of gefitinib within 100 days. Cells that did not acquire resistance within this timeframe (A3B-off H) were excluded from the data set. We characterized these cells and confirmed that the GR cells were markedly resistant to gefitinib (Fig. S4B) and that A3B activity is maintained in GR cells (Fig. S4C).

We next evaluated the prevalence of the EGFR T790M mutation in sensitive and resistant cells (Fig. 3D and Fig. S3B). EGFR T790M mutation was detected in 2 of 5 (40%) A3B-off GR cells (A3B-off GRD and GRE), consistent with previous findings^47,51^. Interestingly, both A3B-off GRD and GRE cells developed gefitinib resistance at early time points, consistent with previous studies suggesting that early gefitinib resistance arises from pre-existing clones with EGFR T790M mutations^47^. In contrast, 5 of 6 (83%) A3B-on GR cells had an EGFR T790M mutation (Fig. 3D). Of these, two A3B-expressing cell lines (A3B-on GRA and GRF) developed gefitinib resistance early, as expected. Unexpectedly, however, three A3B-expressing cell lines (A3B-on GRC, GRD, and GRE) with T790M mutations developed gefitinib resistance at late time-points. This suggests that, in the presence of sustained A3B expression, EGFR T790M mutations are more likely to arise late during the evolution of therapy resistance.

### Integrated genomic analysis of gefitinib-resistant PC9 cells

To gain additional insights into the consequences of APOBEC activity on acquired gefitinib resistance, we performed genomic analyses of control and APOBEC3B-expressing cells, both in gefitinib-sensitive and gefitinib-resistant contexts. Given the role of APOBEC3 enzymes in inducing mutations, we first used whole-exome sequencing to assess the number of mutations present in the coding region of each cell line. We sequenced four control (A3B-off: GRA, GRC, GRD, GRE) and five A3B-expressing (A3B-on: GRA, GRB, GRC, GRD, GRF) gefitinib-resistant cells, along with corresponding gefitinib-sensitive control (A3B-off: A and B) and A3B-expressing (A3B-on: A, B, D, and F) cells. Finally, parental PC9 cells were sequenced as a reference, and mutations observed in parental PC9 cells were filtered out from subsequent analyses (see Methods).

APOBEC3B expression in gefitinib-sensitive cells did not lead to an increase in the number of observed mutations (Fig. 4A; average mutations/exome: A3B-off, 10.5; A3B-on, 9), likely because APOBEC-induced mutations are distributed throughout the genome, and so in a population of cells any individual mutation will be present at very low rates, below the limit of detection by next-generation sequencing^49^. In contrast, the acquisition of gefitinib-resistance was associated with a marked increase in the number of mutations (Fig. 4A; average mutations/exome: A3B-off, 70.3; A3B-on, 90.4). Interestingly, A3B-on GR cells did not have significantly more mutations than A3B-off GR cells, suggesting that the increased number of mutations observed in GR cells was independent of ectopic A3B expression. There are several possible explanations for the increased mutational burden, including other mutagenic processes operative in these cells, the selection for pre-existing subclones with distinct mutations, or the activity of endogenous A3A and A3B induced by gefitinib treatment. Consistent with this latter possibility, both A3B-off and A3B-on GR cells had an increase in C-to-T transitions and C-to-G transversions occurring in a T<u>C</u>W context, which is the APOBEC consensus motif (Supplemental File 1). While the total number of mutations was too low to extract mutational signatures, the presence of these APOBEC consensus mutations in GR cells suggests a contribution of endogenous APOBEC activity to the total mutational burden.

**Figure 4.**
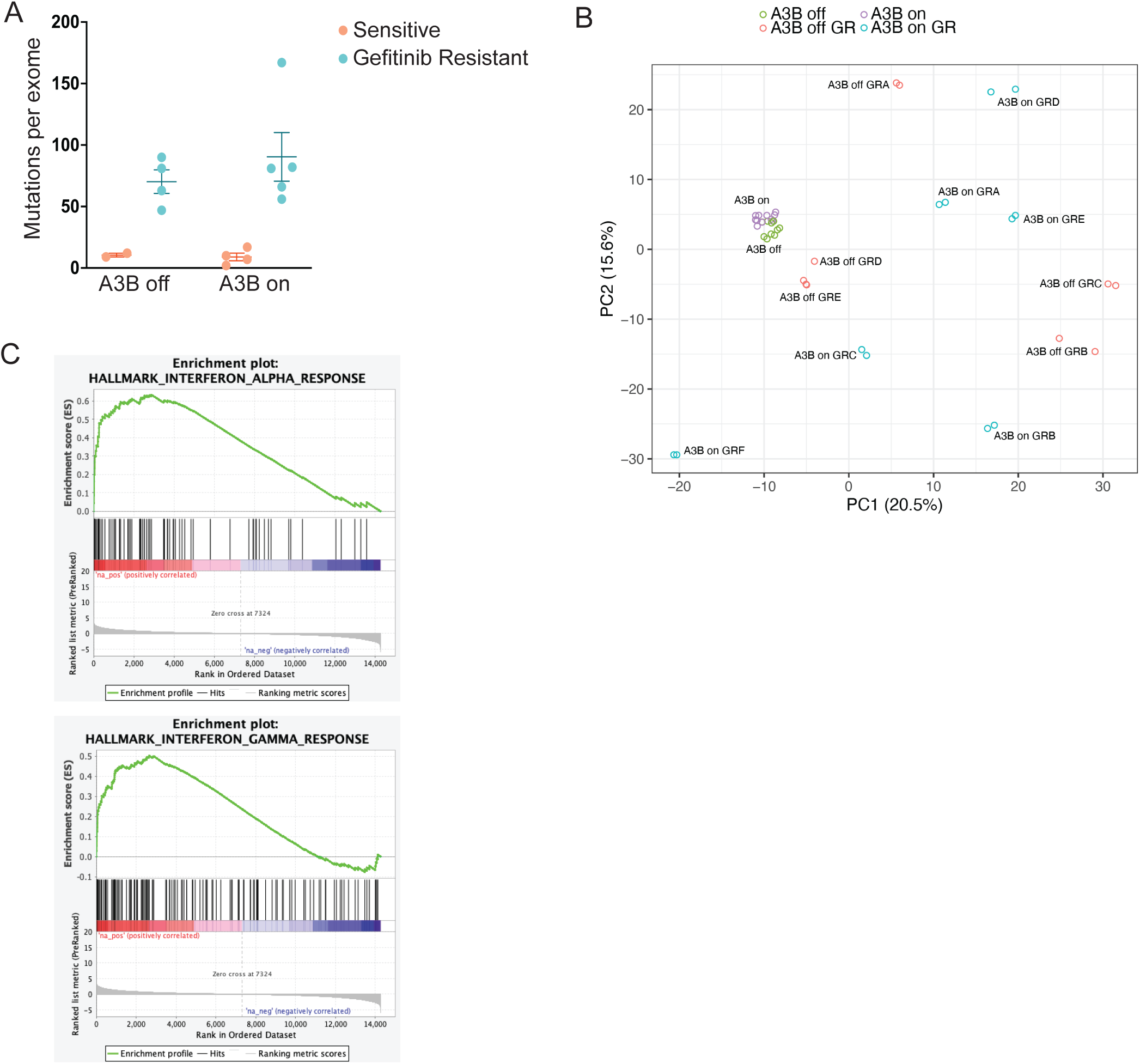
Integrated genomic analysis of gefitinib-resistant PC9 cells. A. Quantification of mutations per exome in A3B off and A3B on gefitinib-resistant cells. Mutations were called as described in Methods. B. Principle component analysis from RNA sequencing data. C. Gene set enrichment analysis comparing A3B-on and A3B-off sensitive PC9 cells.

Examination of mutations specific to GR cells revealed that none of the commonly mutated genes in lung adenocarcinoma, squamous carcinoma, or small-cell cancer was mutated in A3B-off or A3B-on GR cells (data not shown). Similarly, with the exception of EGFR T790M mutations (see above), none of the known genetic mechanisms of resistance to EGFR inhibition was observed in these cells (data not shown). Therefore, we reasoned that non-genetic mechanisms may be responsible for gefitinib resistance in these cells. To address this, we performed RNA sequencing on all A3B-off and A3B-on cells. We first assessed the effect of A3B expression on gene expression in gefitinib-sensitive PC9 cells using RNA-sequencing. Principal components analysis (PCA) of control and A3B-expressing gefitinib-sensitive cells revealed that A3B expression led to modest but consistent changes in gene expression (Fig. 4B). Gene set enrichment analysis showed that A3B expression led to upregulation of pro-inflammatory gene expression programs, including IL6_JAK_STAT3_SIGNALING, INTERFERON_ALPHA_RESPONSE, and INTERFERON_GAMMA_RESPONSE (Fig. 4C and Supplemental File 2).

We next compared the transcriptional profiles of gefitinib-resistant cells with and without A3B expression. A3B- off GR cells with a T790M mutation (A3B-off GRD and GRE) clustered near gefitinib-sensitive cells on a PCA plot, consistent with the notion that these cells have sustained EGFR signaling and very similar gene expression profiles to gefitinib-sensitive cells (Fig. 4B). In contrast, A3B-off GR cells lacking the T790M mutation (A3B- off GRA, GRB, and GRC) were more distant on the PCA plot, suggesting these cells have distinct transcriptional profiles, possibly due to activation of different oncogenic programs in these cells (Fig. 4B). These results are also consistent with previous findings that PC9 cells that develop resistance early are transcriptionally more similar to parental PC9 cells, while PC9 cells that develop EGFR inhibitor resistance at late time-points, via evolution in the DTP state, have distinct gene expression profiles^47^.

We next examined the transcriptional profile of A3B-on gefitinib resistant cells. Interestingly, A3B-on GR cells had gene expression patterns that were distinct from both parental cells and from one another, irrespective of their T790M status (Fig. 4B). This is consistent with the possibility that A3B expression promotes the evolution of resistance during the DTP state, leading to gefitinib-resistant cells with divergent transcriptional programs.

### APOBEC3B-expressing PC9 cells show evidence of squamous cell transdifferentiation during acquired gefitinib resistance

We next examined RNA-seq data for evidence of non-genetic resistance mechanisms in GR cells. Given the relationship between APOBEC mutagenesis and histological transformation, we first asked whether any GR cells had evidence of small-cell or squamous cell transdifferentiation. None of the GR cells exhibited changes in expression of genes encoding the neuroendocrine markers Synaptophysin or Chromagranin, which are expressed in small-cell lung cancer (data not shown). Furthermore, expression of RB1, TP53, and MYC, remain consistent between A3B-off and A3B-on sensitive and resistant cells (Fig. S5B). Loss of both Rb and p53, and amplification of Myc, are associated with small-cell carcinoma histology^29,52,53^. Together, these data suggest that gefitinib resistance in these cells is not associated with transdifferentiation to neuroendocrine, small-cell lung cancer.

We next examined expression of the squamous cell transcription factor p63, which is essential for development of squamous epithelium and highly expressed in squamous cell cancers, including squamous lung cancer^54,55^. RNA-seq data revealed that two A3B-on resistant cell lines – GRB and GRF – had high expression of p63 (Fig. 5A). In contrast, none of the five A3B-off resistant cell lines upregulated p63. qRT-PCR confirmed this finding and showed that expression of both the oncogenic variant of p63, ΔNp63, as well as total p63, were higher in the resistant cells compared to the matched sensitive lines (Fig. 5B). Immunohistochemical staining for ΔNp63 further validated that these GR cells have high p63 expression, both *in vitro* and when grown as xenograft tumors *in vivo* (Fig. 5D – E and S6B).

**Figure 5.**
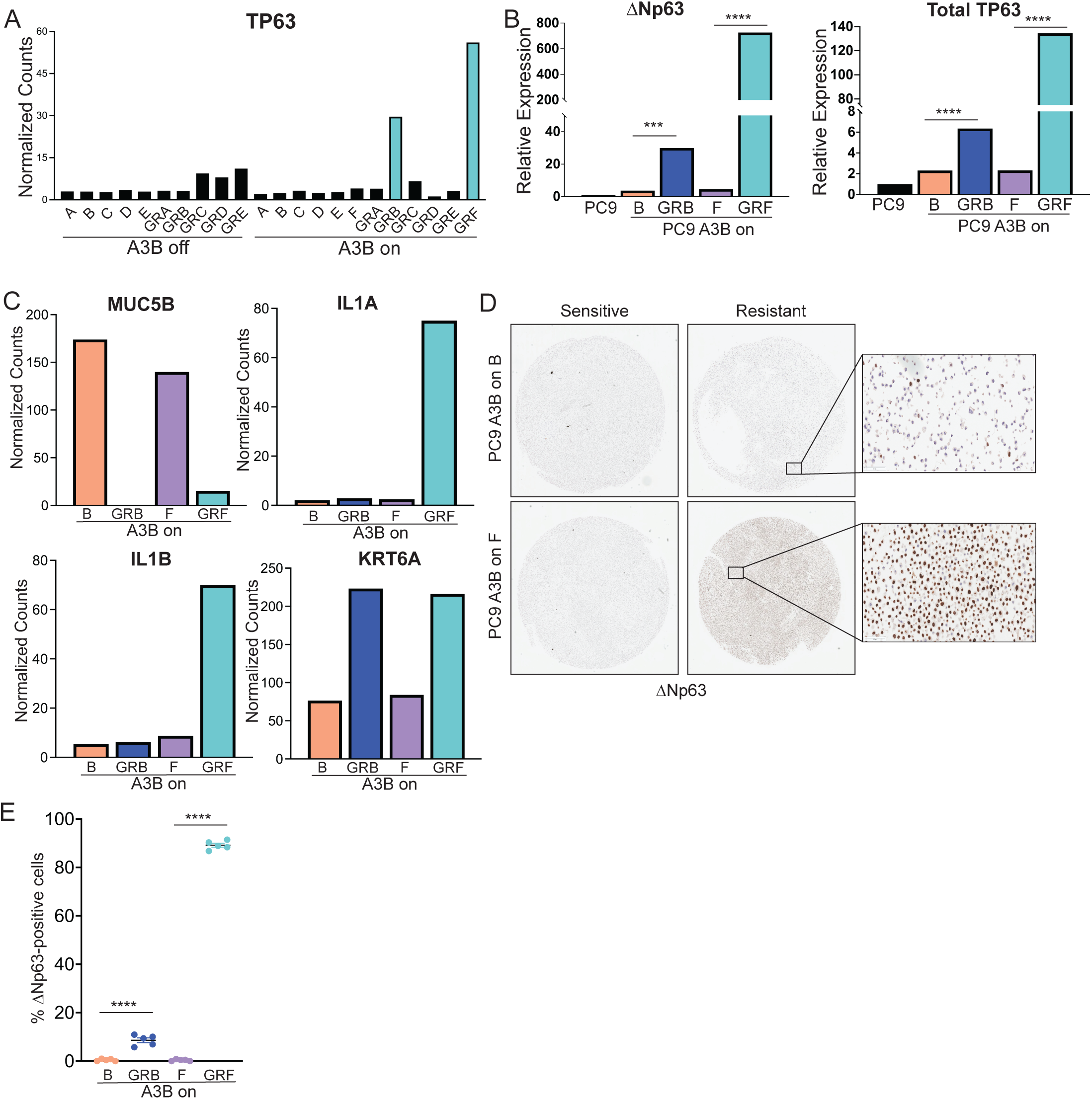
APOBEC3B-expressing PC9 cells show evidence of squamous cell transdifferentiation during acquired gefitinib resistance. A. Normalized gene counts for TP63 from RNA sequencing in sensitive and gefitinib-resistant PC9 cells, with or without A3B expression. B. qRT-PCR analysis showing ΔNp63 and total p63 mRNA expression in sensitive and gefitinib-resistant PC9 A3B-on cells. Parental PC9 are shown as a control. Error bars represent SEM of 3 technical replicates. Unpaired t-test was performed to determine statistical significance. *** indicates p<0.0005 and **** indicates p<0.0001. C. Normalized gene counts for MUC5B, IL1A, IL1B, and KRT6A from RNA sequencing in sensitive and gefitinib-resistant PC9 cells expressing A3B. D. Immunohistochemistry images showing ΔNp63 protein expression in sensitive and resistant PC9 A3B-on cells. Insets are shown at 20X magnification. E. Quantification of ΔNp63-positive cells for sensitive and resistant PC9 A3B-on cells shown in D. Each point represents a field of view at 20X magnification. Error bars represent SEM of five fields of view. Unpaired t-test was performed to determine statistical significance. *** indicates p<0.0005 and **** indicates p<0.0001.

To extend these data, we used qRT-PCR to analyze ΔNp63 expression in 4 additional A3B-off and 5 A3B-on GR lines (see Fig. S4A). We found that one A3B-off (A3B-off GRG) and two A3B-on (A3B-on GRG and GRH) lines exhibited high expression of ΔNp63 (Fig. S6A). In total, 1 of 9 A3B-off and 4 of 11 A3B-on GR cell lines showed upregulation of ΔNp63, suggesting that upregulation of this transcription factor occurs more frequently during acquired resistance to gefitinib in cells with constitutive expression of A3B.

p63 is a known regulator of lineage plasticity in cancer cells^54,56^ and its expression is associated with squamous cell carcinoma transdifferentiation^31^. Consistent with this, we found that the adenocarcinoma marker, *MUC5B*, was downregulated in A3B-on GRB and GRF cells (Fig. 5C). Moreover, SCC marker, *KRT6A,* and p63 target genes, *IL1A* and *IL1B,* were upregulated in A3B-on cells with high ΔNp63 expression (Fig. 5C and S6A). Taken together, these data suggest that A3B-on GRB and GRF cells transdifferentiated to a squamous cell phenotype during the acquisition of gefitinib-resistance. Interestingly, GRF cells have a T790M mutation, while GRB cells do not (Fig. 3D); this mirrors the clinical findings that squamous cell transdifferentiation can occur in the presence or absence of T790M^57^.

### p63 knockout reduces inflammatory gene expression and sensitizes PC9 GR cells to EGFR inhibition

To evaluate the function of ΔNp63 in gefitinib-resistant cells we used A3B-on GRF cells, since these expressed high levels of ΔNp63. We first measured sensitivity to osimertinib, a third-generation EGFR inhibitor effective against the T790M mutation. As controls we used two gefitinib-sensitive lines, A3B-on F and E, and the gefitinib-resistant line A3B-on GRE, which has an EGFR T790M mutation but does not express ΔNp63. As expected, A3B-on GRE cells (IC50: 12.66 nM) were sensitive to osimertinib, with IC50 values similar to the gefitinib-sensitive lines (A3B-on F: 17.48 nM, A3B-on E: 14.11 nM). In contrast, A3B-on GRF cells (IC50: 81.97 nM), which express ΔNp63, were more resistant to osimertinib treatment (Fig. 6A). To explore the molecular basis for this difference, we tested whether osimertinib effectively inhibited EGFR signaling in these cells. Whereas treatment with 100 nM osimertinib completely abolished EGFR phosphorylation in gefitinib-sensitive and A3B- on GRE cells, this dose of drug only partially inhibited EGFR phosphorylation in A3B-on GRF (Fig. 6B).

**Figure 6.**
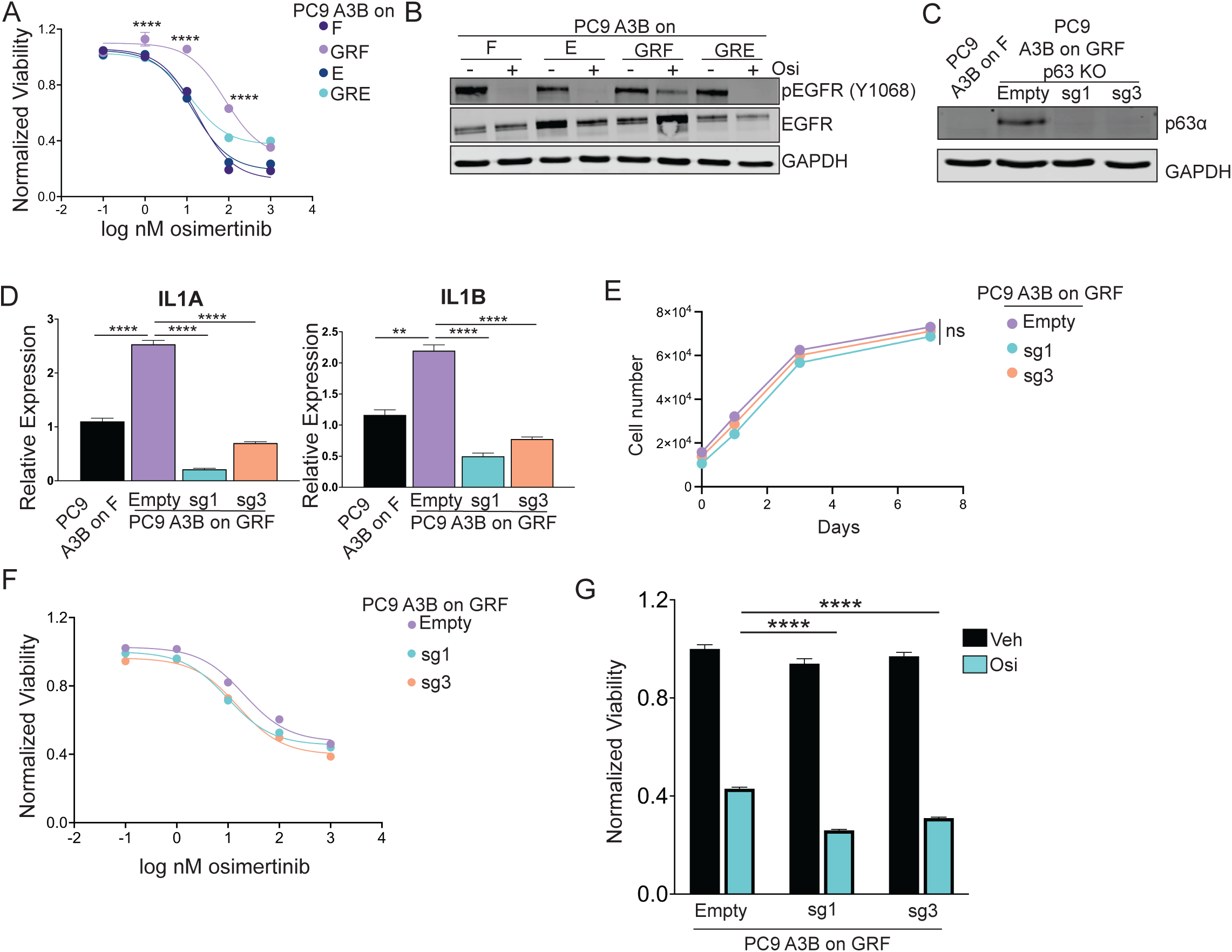
p63 knockout reduces inflammatory gene expression and sensitizes PC9 GR cells to EGFR inhibition. A. Dose-response curve to osimertinib in sensitive and gefitinib-resistant PC9 cells expressing A3B. Viability was measured using CellTiter Glo following 3 days of drug treatment. Two-way ANOVA with Tukey’s multiple comparisons test was used to determine statistical significance at each dose. **** indicates p<0.0001 when comparing PC9 A3B-on GRF and GRE cells at the indicated dose. B. Western blot for pEGFR (Y1068) and total EGFR in sensitive and gefitinib-resistant PC9 cells expressing A3B. Cells were treated with osimertinib for 24 hours. C. Western blot for p63α in PC9 A3B-on GRF cells expressing an empty vector or gRNAs targeting p63 (sg1 and sg3). PC9 A3B-on F cells are shown as a control. D. qRT-PCR analysis showing IL1A and IL1B expression in PC9 A3B-on GRF cells expressing an empty vector or gRNAs targeting p63 (sg1 and sg3). PC9 A3B-on F cells are shown as a control. Error bars represent SEM of three technical replicates. Unpaired t-test was performed to determine statistical significance. ** inidicates p<0.005 and **** indicates p<0.0001. E. In vitro growth curves of PC9 A3B-on GRF cells with or without p63 knockout. Two-way ANOVA was performed to determine statistical significance. ns = not significant. F. Dose-response curve to osimertinib in PC9 A3B-on GRF cells expressing an empty vector or gRNAs targeting p63 (sg1 and sg3). Viability was measured using CellTiter Glo following 3 days of drug treatment. Two-way ANOVA was performed and determined a significant p63 status × dose interaction with a p value of 0.02. G. Viability of PC9 A3B-on GRF cells expressing an empty vector or gRNAs targeting p63 (sg1 and sg3) following treatment with osimertinib for 6 days. A two-way ANOVA with Dunnett’s test was performed to determine statistical significance. **** indicates p<0.00001.

To test whether high expression of ΔNp63 directly promotes osimertinib resistance in these cells, we knocked out the *TP63* gene using CRISPR-Cas9 (Fig. 6C). Expression of the ΔNp63 target genes *IL1A* and *IL1B* was reduced in p63-knockout cells, confirming functional knockout (Fig. 6D). While p63 knockout did not affect the growth of A3B-on GRF cells *in vitro* (Fig. 6E), its knockout sensitized these cells to osimertinib treatment over 3 and 6 days (Fig. 6F – G). A3B-on GRF cells expressing a control vector had an IC50 of 20.40 nM, which was lower than the IC50 of parental A3B-on GRF cells (cf. Fig. 6A). Nonetheless, A3B-on GRF p63 KO 1 and KO 3 cells had IC50s of 9.89 nM and 15.60 nM, respectively. These findings show that high expression of ΔNp63 in gefitinib-resistant cells with a T790M mutation promotes resistance to third-generation EGFR inhibitors in these cells.

## DISCUSSION

TKIs that target EGFR are the current standard of care for patients with EGFR-mutant lung adenocarcinomas. Although these treatments are initially effective, resistance to TKIs occurs within months^16–18^. Understanding the mechanisms that can lead to acquired drug resistance can provide better insight into how to treat this subset of patients.

In the current study, we show that targeted therapies against EGFR induce APOBEC activity in an EGFR-mutant non-small cell lung cancer cell line. EGFR inhibition leads to a transient increase in A3A and A3B expression and APOBEC deaminase activity. This mirrors clinical data that shows an increase in the APOBEC mutational signature in tumors from patients who have acquired resistance to an EGFR inhibitor, especially those with tumors that have undergone histological transformation^32–34^. While sustained APOBEC activity does not accelerate acquired therapy resistance, A3B expression alters the evolutionary path that PC9 cells take to become gefitinib-resistant. Specifically, A3B expression is associated with the late acquisition of T790M mutations during the DTP state. Consistent with this, knockout of A3A and A3B impairs the survival of DTPs. Taken together, these data support a model where induction of APOBEC activity promotes DTP survival, thereby facilitating the on-going evolution of drug-tolerant persister cells. Importantly, these results corroborate recent findings from other groups demonstrating a role for therapy-induced APOBEC3 expression in DTP survival^35,36^.

A subset of gefitinib-resistant cells displays ΔNp63 upregulation and evidence of squamous cell transdifferentiation, and importantly ΔNp63 upregulation was more common in GR cells expressing A3B than in control cells. This suggests that APOBEC activity may be functionally linked to this form of histological transformation. p63 is a p53-family member known for its role in maintaining stemness of epithelial cells^56,58^. Work from other labs has demonstrated that p63 amplification and, in turn, transcription of its target genes promotes cancer development and progression^54,55,59^. Our findings suggest that ΔNp63 contributes to TKI resistance in squamous transdifferentiated lung cancer cells, since knockout of p63 sensitized these cells to third-generation EGFR inhibitors, like osimertinib. These data may explain the clinical observation that lung cancers exhibiting transdifferentiation have a particularly poor prognosis. Furthermore, these findings may suggest avenues for future therapies. Using ΔNp63 expression as a biomarker, for example, may predict whether or not a patient will respond to third-generation EGFR inhibitors. Therapies against ΔNp63 targets, such as IL1α/β, may also prove useful in the clinical setting. In conclusion, our findings provide mechanistic insights into the link between APOBEC mutagenesis, TKI resistance, and histological transformation in EGFR-mutant lung cancer. More broadly, our work identifies a role for APOBEC3 induction in promoting the survival of DTPs following oncogene-targeted therapies. Future work will define whether APOBEC3 induction is a common response to diverse targeted therapies, and whether APOBEC-dependent survival of DTPs can promote therapy resistance across different tumor types.

## ACKNOWLEDGEMENTS

We thank Dr. John Maciejowski (MSKCC) for providing the A3B antibody. This work was funded by the National Cancer Institute under award R01CA208042 (to J.V.A.), the American Cancer Society under award 132556-RSG-18-130-CCG (to J.V.A.), an HHMI Gilliam Fellowship (to N.M.G.G), and by startup funds from the Fred Hutchinson Cancer Center (to J.V.A.). This research was supported by the following shared resources of the Fred Hutch/University of Washington/Seattle Children’s Cancer Consortium (P30 CA015704): Genomics & Bioinformatics Shared Resource, RRID:SCR_022606; Experimental Histopathology Shared Resource, RRID:SCR_022612; Comparative Medicine Shared Resource, RRID:SCR_022610; and Flow Cytometry Shared Resource, RRID:SCR_022613.

