## Supplemental Figures for "APOBEC3 activity promotes the survival and evolution of drug-tolerant persister cells during acquired resistance to EGFR inhibitors in lung cancer"

Figure S1 (Related to Figure 1)

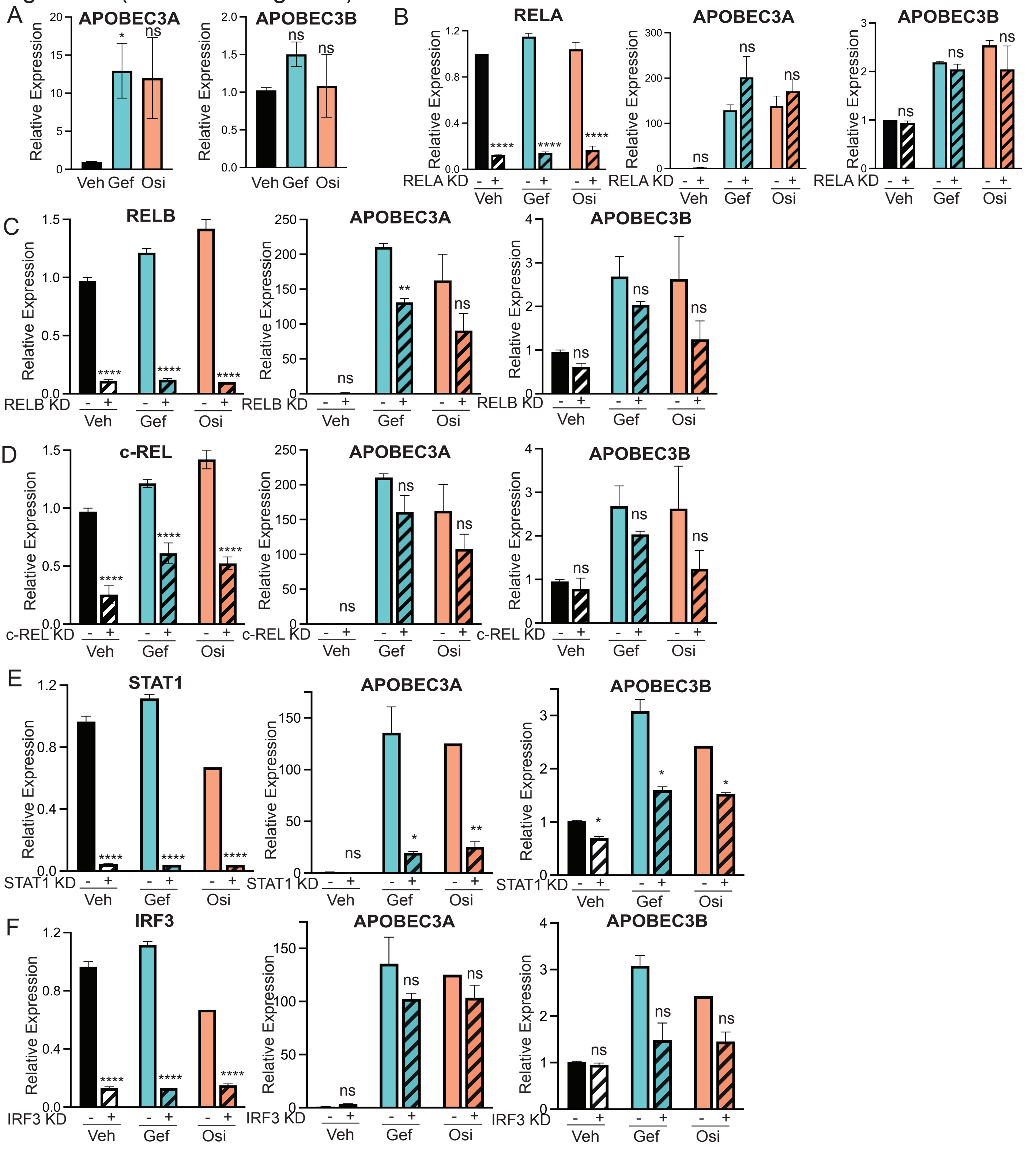

**Figure S1 (Related to Figure 1). APOBEC expression is partially regulated by NFkB and STAT1.**

A. qRT-PCR analysis showing A3A and A3B expression in HCC827 cells following treatment with gefitinib or osimertinib for 24 hours. Error bars represent SEM of two biological replicates. Significance relative to the vehicle was determined using unpaired t-test. * indicates $p<0.05$ and ns = not significant.

B. qRT-PCR analysis showing RELA, A3A, and A3B expression in PC9 cells following RELA knockdown and treatment with gefitinib or osimertinib. Data represents a 48-hour siRNA transfection and 24-hour drug treatment. Error bars represent SEM of 3 technical replicates. Unpaired t-test relative to non-targeting ( - ) was used to determine statistical significance. **** indicates $p<0.0001$ and ns = not significant.

C. qRT-PCR analysis showing RELB, A3A, and A3B expression in PC9 cells following RELB knockdown and treatment with gefitinib or osimertinib. Data represents a 48-hour siRNA transfection and 24-hour drug treatment. Error bars represent SEM of 3 technical replicates. Unpaired t-test relative to non-targeting ( - ) was used to determine statistical significance. ** indicates $p<0.005$, **** indicates $p<0.0001$, and ns = not significant.

D. qRT-PCR analysis showing c-REL, A3A, and A3B expression in PC9 cells following c-REL knockdown and treatment with gefitinib or osimertinib. Data represents a 48-hour siRNA transfection and 24-hour drug treatment. Error bars represent SEM of 3 technical replicates. Unpaired t-test relative to non-targeting ( - ) was used to determine statistical significance. **** indicates $p<0.0001$, and ns = not significant.

E. qRT-PCR analysis showing STAT1, A3A, and A3B expression in PC9 cells following STAT1 knockdown and treatment with gefitinib or osimertinib. Data represents a 48-hour siRNA transfection and 24-hour drug treatment. Error bars represent SEM of 3 technical replicates. Unpaired t-test relative to non-targeting ( - ) was used to determine statistical significance. * indicates $p<0.05$, ** indicates $p<0.005$, **** indicates $p<0.0001$, and ns = not significant.

F. qRT-PCR analysis showing IRF3, A3A, and A3B expression in PC9 cells following IRF3 knockdown and treatment with gefitinib or osimertinib. Data represents a 48-hour siRNA transfection and 24-hour drug treatment. Error bars represent SEM of 3 technical replicates. Unpaired t-test relative to non-targeting ( - ) was used to determine statistical significance. **** indicates $p<0.0001$, and ns = not significant.

Figure S2 (Related to Figure 1)

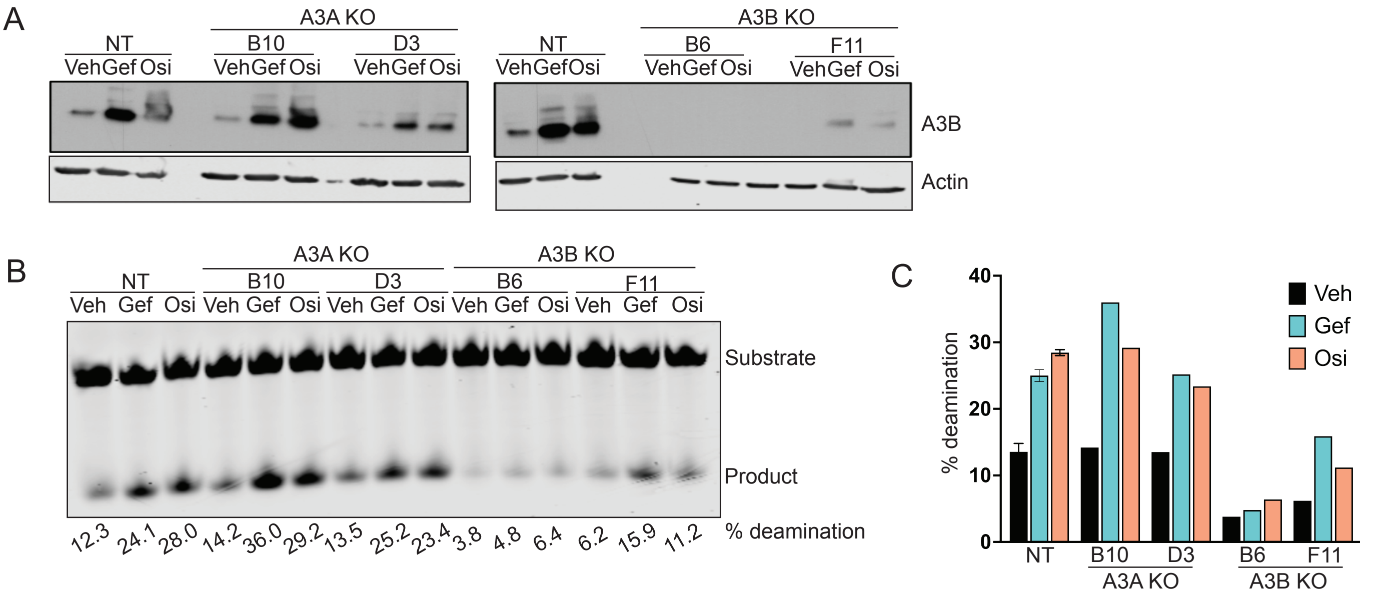


**Figure S2 (Related to Figure 1). EGFR inhibitor-induced APOBEC activity is attenuated following single knockouts of A3A and A3B.**

A. Western blot for A3B protein in PC9 cells expressing non-targeting gRNAs (NT) or gRNAs targeting A3A (A3A KO) or A3B (A3B KO). A pooled population is shown for NT cells, and two clones each are shown for A3A KO and A3B KO. Cells were treated with gefitinib or osimertinib for 24 hours.

B. In vitro deaminase activity of PC9 cells expressing non-targeting gRNAs (NT) or gRNAs targeting A3A (A3A KO) or A3B (A3B KO). A pooled population is shown for NT cells, and two clones each are shown for A3A KO and A3B KO. Cells were treated with gefitinib or osimertinib for 24 hours and % deamination was calculated as described in Methods.

C. Quantification of % deamination shown in B. % deamination was calculated as described in Methods.

Figure S3 (Related to Figure 3)

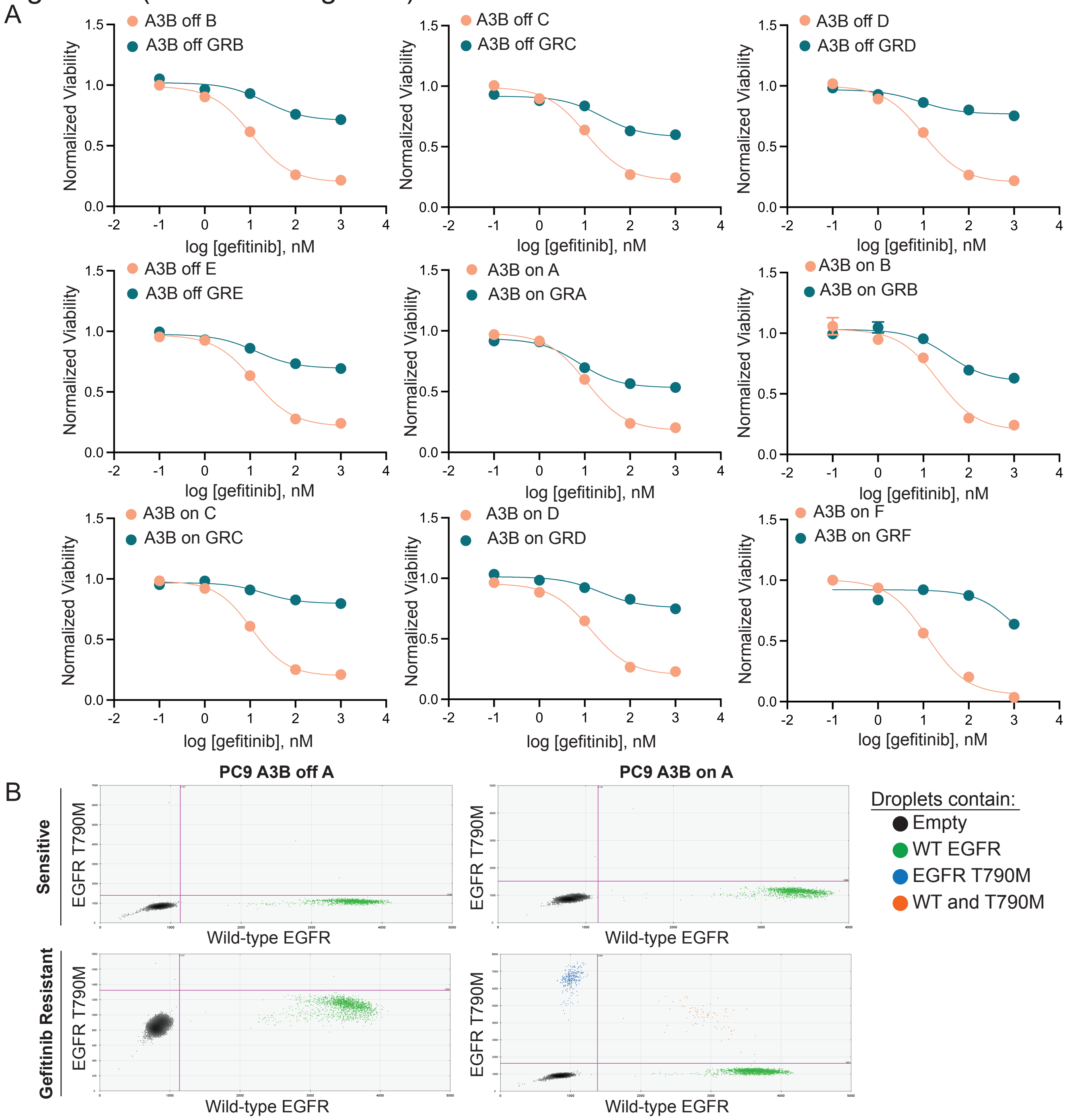


**Figure S3 (Related to Figure 3). PC9 cells expressing APOBEC3B maintain high APOBEC activity following resistance to gefitinib.**

A. Dose-response curves to gefitinib for sensitive and resistant A3B-off and A3B-on PC9 cells.

B. Representative images of ddPCR analysis of sensitive or gefitinib-resistant PC9 cells, with or without A3B expression.

Figure S4 (Related to Figure 3)

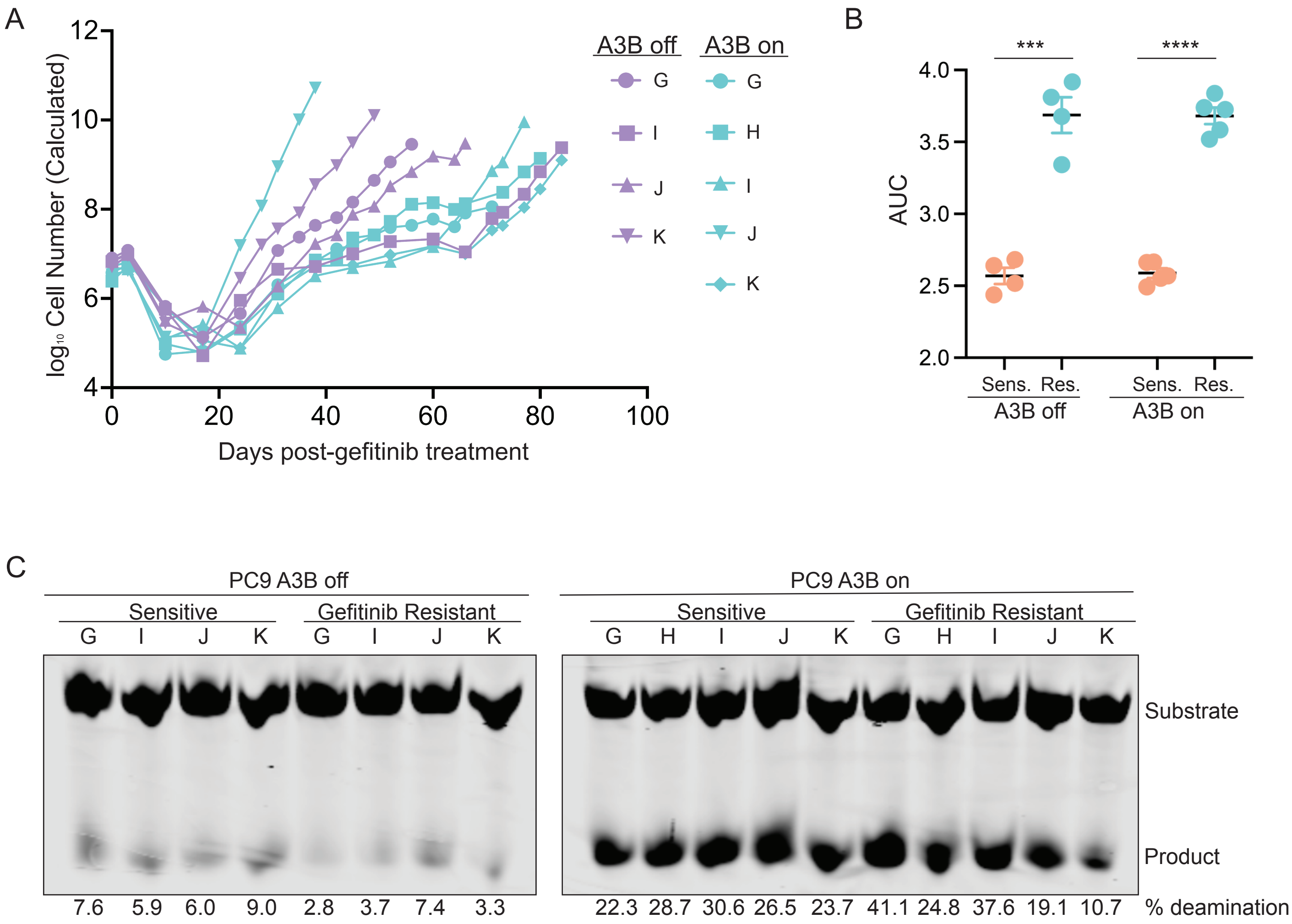


**Figure S4 (Related to Figure 3). Developing resistance to gefitinib in A3B-off and A3B-on PC9 cells.**

A. Kinetics of evolution of gefitinib resistance in PC9 cells with or without A3B expression.

B. Quantification of area under the curve (AUC) of dose-response curves to gefitinib for sensitive and resistant A3B-off and A3B-on PC9 cells. Unpaired t-test was performed to determine statistical significance. *** indicates $p<0.0005$.**** indicates $p<0.0001$.

C. In vitro deaminase activity of sensitive or gefitinib resistant PC9 cells with or without A3B expression. % deamination was calculated as describes in Methods.

Figure S5 (Related to Figure 5)

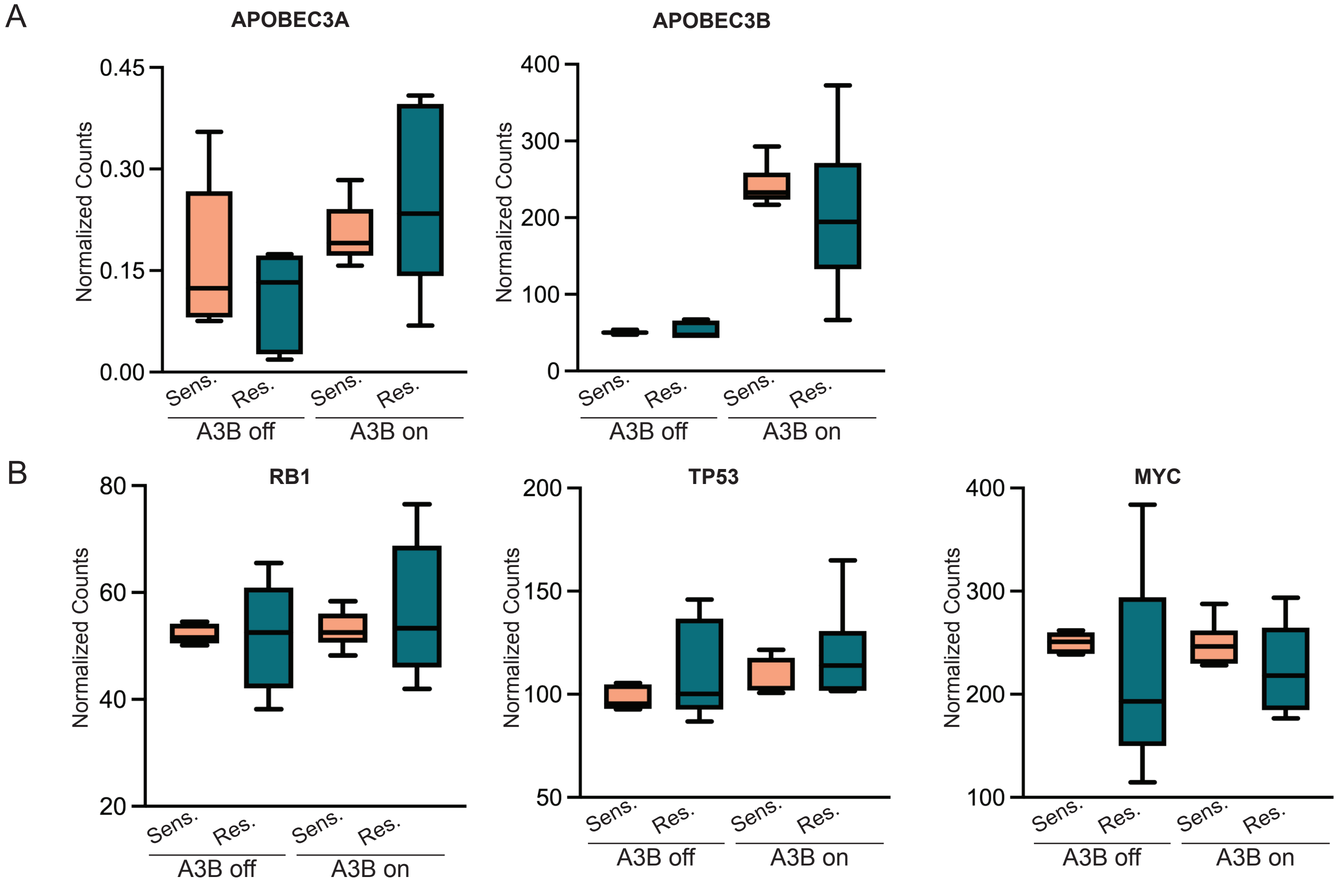


**Figure S5 (Related to Figure 5). Gefitinib-resistant PC9 cells do not express markers of small-cell lung cancer transdifferentiation.**

A. Normalized counts for APOBEC3 expression from RNA-sequencing data analysis of sensitive and gefitinib-resistant PC9 cells, with or without A3B expression.

B. Normalized counts for small-cell lung cancer markers, RB1, TP53, and MYC, from RNA-sequencing data analysis of sensitive and gefitinib-resistant PC9 cells, with or without A3B expression.

Figure S6 (Related to Figure 5)

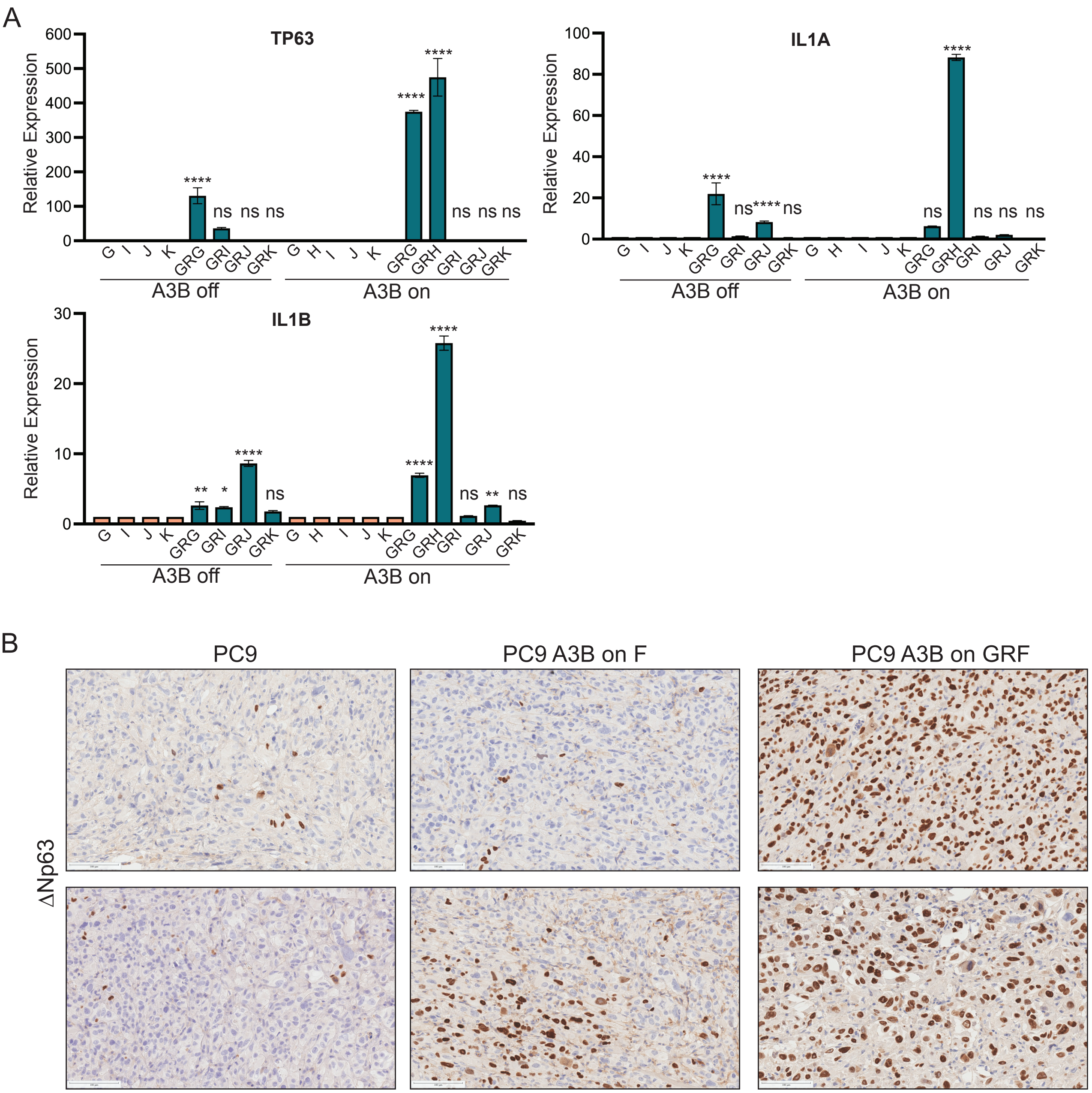


**Figure S6 (Related to Figure 5). p63 and its target genes are highly expressed in A3B-on gefitinib-resistant PC9 cells.**

A. qRT-PCR analysis showing TP63, IL1A, and IL1B expression in sensitive and gefitinib-resistant PC9 cells, with or without A3B expression. Error bars represent three technical replicates. One-way ANOVA with Sidak's multiple comparisons test was used to determine statistical significance relative to each sensitive control. * indicates $p<0.05$, ** indicates $p<0.005$, **** indicates $p<0.0001$, and ns = not significant.

B. Immunohistochemistry images showing ΔNp63 protein expression in parental PC9, PC9 A3B-on F, and PC9 A3B-on GRF xenograft tumors. Images are shown at 20X magnification.
